# Model-optimized stimulus distortions for adaptive estimation of individual sensory representations

**DOI:** 10.64898/2026.07.02.736141

**Authors:** Josue Casco-Rodriguez, Fangfang Hong, David H. Brainard, Jenelle Feather, David Lipshutz

## Abstract

Representations of the same physical stimulus vary between individuals. Characterizing individual differences has practical implications, but is challenging because these representations are not directly observable. Given a model of how representations vary within a population, we propose a Bayesian adaptive procedure for estimating an individual observer’s representation from a series of targeted perceptual discrimination judgments. A key component of our approach is using Fisher information to identify stimulus distortions that efficiently differentiate observers in the population. As a proof of concept, we focus on individual differences in color perception and simulate observers with cone fundamentals drawn from an individual colorimetric observer model. We demonstrate that our approach can recover key aspects of a sampled observer’s cone fundamentals using simulated three-alternative forced-choice oddity judgments with approximately 500 trials, corresponding to an experimental duration of approximately one hour. Our Bayesian adaptive framework provides a promising and generalizable approach to efficiently link behavioral measurements to individual differences in sensory representations.

## 1 Introduction

Quantifying perceptual differences between individual observers is a core challenge across many neuroscience and cognitive science domains (Hirsh & Watson, 1996; Drayna, 2005; Keller, Zhuang, Chi, Vosshall, & Matsunami, 2007; Hasin-Brumshtein, Lancet, & Olender, 2009; Wilmer, 2008; Mainland et al., 2014; Mollon, Bosten, Peterzell, & Webster, 2017). While some variations can be attributed to well-characterized clinical deficits, such as color vision deficiencies or hearing impairment, other subtle differences are harder to detect. For example, images such as “The Dress” demonstrate that healthy observers can disagree dramatically about the appearance of the same stimulus (Brainard & Hurlbert, 2015). Such examples suggest that even more subtle and idiosyncratic differences in perception are ubiquitous across our everyday experiences. However, measuring these differences is challenging because the underlying sensory representations mediating the differences are often not directly observable.

Here we develop a Bayesian framework to infer an observer’s underlying sensory representation based on a series of perceptual discrimination judgments. Given a parametric model of individual variability of sensory representations, we first identify stimulus distortions where observers are expected to maximally disagree. Building on the work of Zhou, Chun, Subramanian, and Simoncelli (2024) and Feather, Lipshutz, Harvey, Williams, and Simoncelli (2025), we approximate observers’ perceptual sensitivities using Fisher information, which reduces the selection of an informative stimulus distortion to a standard eigenvalue problem, and thus avoids the combinatorial explosion of simulating each observer’s response to every possible stimulus distortion. We embed this procedure in a Bayesian adaptive loop: on each iteration, we select a stimulus distortion based on the current distribution over individuals, measure the observer’s perceptual discrimination judgments, and update the posterior.

We provide a proof of concept for this approach in the domain of color vision, where individual variation in physiological properties of the retina can lead to measurable differences in perceptual appearance even among normal-vision observers. Specifically, differences in pre-receptoral absorption, cone photopigment densities, and cone photopigment spectral absorbances give rise to individual differences in cone fundamentals (CFs). As a result, the same physical stimulus can produce different cone excitations for different observers, and therefore potentially different perceptual appearances (Stiles & Burch, 1959; MacLeod & Webster, 1983; Asano, Fairchild, Blondé, & Morvan, 2016; Mollon et al., 2017). Understanding this variation has practical implications for color reproduction, where individual color vision profiles could inform the design of wide-gamut displays and color-rendering algorithms, either by reducing perceived color differences across observers or by tailoring color reproduction to a specific observer (Long & Fairchild, 2015; Stockman & Rider, 2025; Stockman, Luo, Shi, Song, & Rider, 2025).

More concretely, we apply our framework to the problem of estimating individual CFs from simulated color discrimination judgments. Using the individual colorimetric observer model of Asano, Fairchild, and Blondé (2016), we compute the cone responses for colors rendered on a 16-channel display (Oh et al., 2022), and simulate observer performance on a three-alternative forced choice (3AFC) color-discrimination task following (Hong et al., 2025; see also, Krauskopf & Gegenfurtner, 1992; Danilova & Mollon, 2025). Through simulated experiments, we show that the procedure accurately identifies key aspects of individual observers’ CFs from a modest number of discrimination measurements. This efficiency is important for practical applications: a brief viewer-specific calibration could provide the information needed to implement personalized color-rendering algorithms. The remainder of this paper is organized as follows: we first present the general framework, which applies to any individual observer model that maps sensory stimuli to individual representations; we then apply the framework to individual colorimetry, describing the observer model, simulated display, and simulated experimental design; and finally, we evaluate how accurately the adaptive procedure recovers individual CFs in simulation.

## 2 Methods

### 2.1 Modeling individual variability in sensory representations

We first present a general framework for characterizing the individual variability in sensory representations. Given a stimulus ***s***, we assume that each individual’s internal responses are stochastic and described by a conditional density. Specifically, we model these conditional densities using a parameterized family *q*_***θ***_ (***r*** | ***s***), where ***s*** is a stimulus vector, ***r*** is the stochastic response vector (e.g., noisy neural responses), and ***θ*** is a vector that parametrizes sensory representations across a population. That is, for each individual, we assume there exists ***θ*** such that *q*_***θ***_ (***r*** | ***s***) well approximates their true stochastic stimulus-response relationship. We further assume a prior distribution *π*(***θ***) over parameters that captures how individuals are distributed within the population. Figure 1A illustrates a toy example of a multimodal prior distribution over a scalar parameter *θ* . Because each parameter ***θ*** corresponds to one sensory representation, drawing ***θ*** from *π*(***θ***) corresponds to sampling a representation from the population, weighted by how common it is. Although *π*(***θ***) is technically a distribution over *parameters*, throughout this work we often refer to it as a distribution over *observers*.

**Figure 1:**
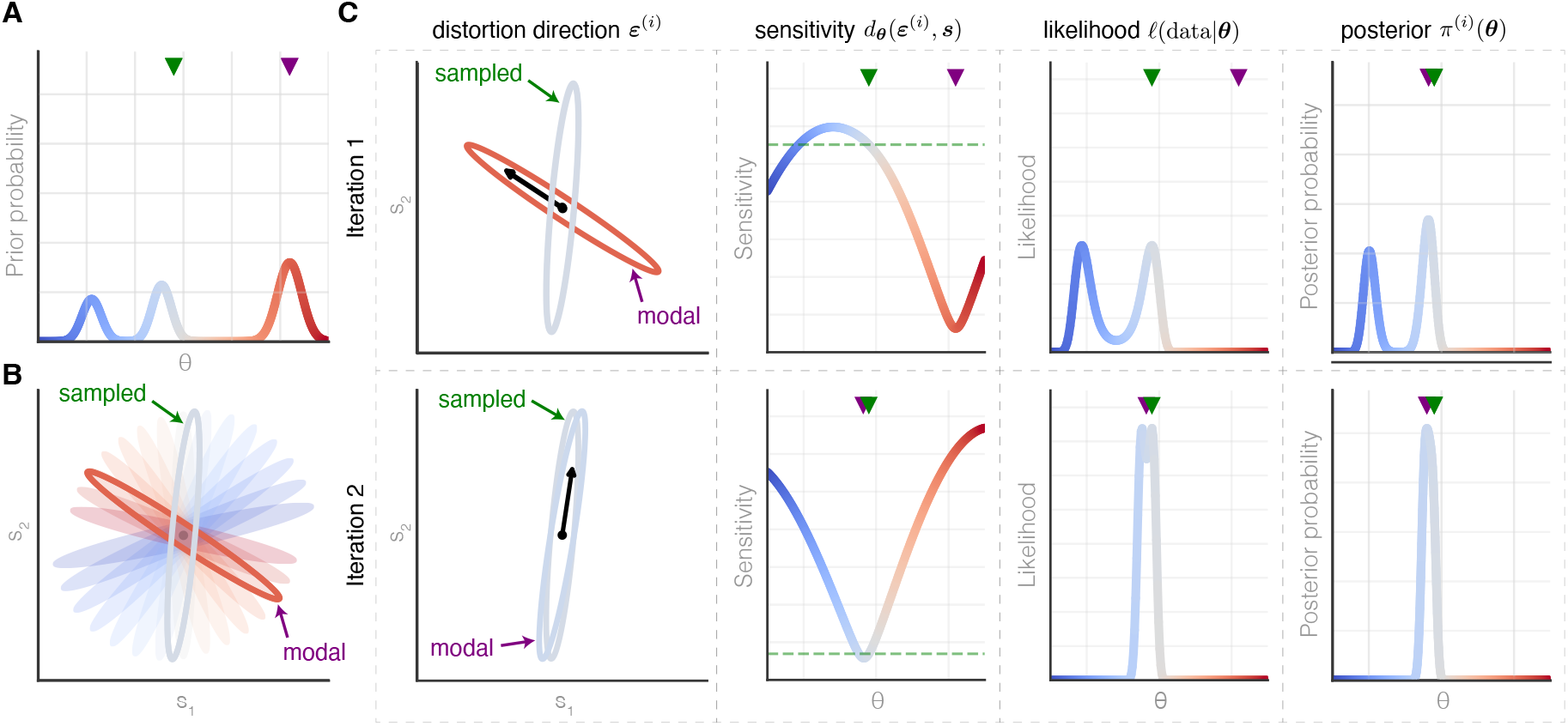
Schematic of our adaptive estimation method. **A.** Prior distribution *π*(*θ* ) over observers. The sampled observer (green triangle) is the unknown observer being estimated. The modal observer (purple triangle) is the current best guess. **B**. Iso-threshold ellipses{ ***u*** : *d*_***θ***_ (***u***; ***s***) = 1} at a base stimulus ***s*** (shown by filled circle). The variation in orientation of the ellipses illustrates differences in perceptual sensitivities across observers. **C**. Bayesian adaptive estimation procedure over 2 iterations. Left column: model-optimized distortion direction ***ε***^(*i*)^ (black arrow) points in a direction along which the average observer is sensitive relative to the modal observer. Center-left column: sensitivity *d*_***θ***_ (***ε***^(*i*)^; ***s***) of each observer along ***ε***^(*i*)^. The sensitivity is minimized around the modal observer. The sampled observer’s sensitivity is indicated with a green horizontal dashed line. Center-right column: likelihood of each observer *ℓ*(*θ* | data) given the measured sensitivity. The likelihood peaks around the sampled observer. Right column: posterior distribution *π*^(*i*)^(***θ***). In iteration 1 (top row), the sampled observer is sensitive to ***ε***^(1)^, so the modal observer is unlikely and the posterior *π*^(1)^(***θ***) shifts away from the modal observer. In iteration 2 (bottom row), the sampled observer is insensitive to ***ε***^(2)^, confirming the modal observer as a good hypothesis, so *π*^(2)^(***θ***) concentrates around the modal observer.

Given an observer randomly sampled from the population, i.e. ***θ*** ∼ *π*(***θ***), we seek an efficient method for estimating ***θ*** based on a series of perceptual discrimination judgments. Consider a base stimulus ***s*** and a distorted stimulus ***s*** + Δ***ε***, where ***ε*** is a unit-norm distortion direction and the scalar Δ (which can be positive or negative) controls the distortion weight. An individual’s ability to discriminate the distorted stimulus from the base stimulus is hypothesized to depend on the overlap between *q*_***θ***_ (***r*** | ***s***) and *q*_***θ***_ (***r*** | ***s*** + Δ***ε***), with less overlap corresponding to smaller discrimination thresholds (Green & Swets, 1966). Discrimination thresholds at a stimulus ***s*** thus reveal the local structure of *q*_***θ***_ (***r*** | ***s***), which can in turn be used to estimate ***θ***.

How should the distortion direction ***ε*** be chosen so that the resulting threshold measurements are maximally informative about ***θ***? One approach is to directly simulate observer responses to estimate their discrimination thresholds. However, this is computationally prohibitive as it requires estimating a discrimination threshold for every candidate distortion and every candidate observer, with each threshold requiring a large number of simulated trials. As an alternative, we estimate perceptual sensitivity using a local measure that can be computed directly from the observer model.

### 2.2 Quantifying local perceptual sensitivity using Fisher information

We predict each model observer’s perceptual sensitivity using the Fisher information matrix (FIM) (Fisher, 1925), which has been used to connect neural representations to perceptual discrimination (Paradiso, 1988; Brunel & Nadal, 1998; Seriès, Stocker, & Simoncelli, 2009; Ganguli & Simoncelli, 2014; Wei & Stocker, 2016; Berardino, Laparra, Ballé, & Simoncelli, 2017). The FIM captures how distinguishable nearby stimuli are to a given observer who efficiently uses the neural representation: given a base stimulus ***s*** and a distortion direction ***ε***, the KL divergence between the observer’s response distribution at ***s*** and ***s*** + Δ***ε*** is approximately 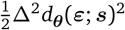, where

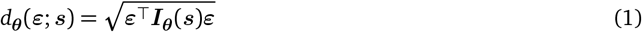

is the *sensitivity* of the representation *q*_***θ***_ at stimulus ***s*** to distortions in the direction ***ε***, and ***I***_***θ***_ (***s***) is the FIM defined by

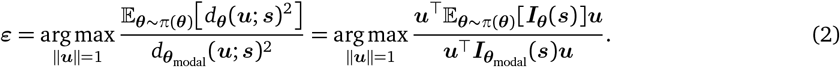

We assume that an observer’s discrimination threshold along distortion direction ***ε*** at base stimulus ***s***—i.e., the smallest |Δ| such that the observer can reliably discriminate between ***s*** and ***s*** + Δ***ε***—is inversely proportional to the scaled sensitivity |Δ| *d*_***θ***_ (***ε***; ***s***): a larger sensitivity means the observer’s internal representation changes more with the distortion relative to the variability of the internal representation, yielding a lower threshold. Under this assumption, the observer’s iso-threshold ellipsoid at ***s***, *T*_***s***_(***θ***) = {***s*** + *d*_***θ***_ (***u***; ***s***)^−1^***u*** : ∥***u***∥ = 1}, is the set of distorted stimuli that are perceptually equidistant to the base stimulus. Differences in sensitivities across observers correspond to differences in ellipse shape, size, and orientation. Figure 1B illustrates these ellipses for a two-dimensional stimulus space, with observers drawn from a prior over a scalar parameter ***θ***. The goal of the next section is to identify distortion directions that are maximally informative about where a given observer falls within this family of ellipses.

### 2.3 Model-optimized stimulus distortions for differentiating visual representations

We seek distortion directions that reveal differences within the family of stimulus representations {*q*_***θ***_ (***r*** | ***s***) }. To build intuition for our approach, first consider the case of two observers: ***θ*** ∈ {*A, B*} . A natural choice is to find a distortion direction that maximizes observer *A*’s sensitivity while minimizing observer *B*’s (Zhou et al., 2024), which is conceptually similar to methods that maximize disagreement between models (Wang & Simoncelli, 2008; Golan, Guo, & Kriegeskorte, 2022). Specifically, a distortion is chosen to extremize the sensitivity ratio:

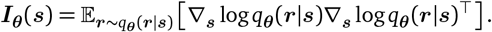

where the optimization is over a unit distortion direction ***u***. From Equation 1, this amounts to optimizing a generalized Rayleigh quotient, and the optimal distortion corresponds to the principal eigenvector of the generalized eigenvalue problem ***I***_*A*_(***s***)***ε*** = *λ****I***_*B*_ (***s***)***ε***, which can be efficiently computed using standard numerical methods (Golub & Van Loan, 2013). Feather et al. (2025) extended this approach to *N* > 2 observers by optimizing for distortions that simultaneously maximize all pairwise sensitivity ratios. However, this pairwise optimization approach does not exploit the smooth structure of the observer population: when observers vary continuously in parameter space, nearby observers have similar FIMs.

We propose an alternative approach that anchors on the *modal* observer (i.e., the most likely observer given the data) and selects a distortion direction to which the modal observer is insensitive relative to the population average. Specifically, letting ***θ***_modal_ = arg max_***θ***_ *π*(***θ***) denote the modal observer, we select the distortion that maximizes the ratio of the expected sensitivity to the modal observer’s sensitivity:

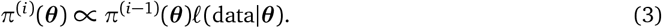

A large ratio indicates a direction ***ε*** that is highly visible to much of the population but not to the modal observer, making the observer’s response useful in either case: if the observer is sensitive to ***ε*** then the posterior shifts away from the modal observer (Figure 1C, Iteration 1), whereas if the observer is insensitive then the posterior further concentrates on the modal observer (Figure 1C, Iteration 2). Equation 2 is also a generalized Rayleigh quotient and thus can also be solved efficiently and incorporated into a closed-loop procedure. In the adaptive procedure described below, in the *i*th iteration, *π*(***θ***) is replaced by the current posterior *π*^(*i*−1)^ with the convention that *π*^(0)^ corresponds to the prior *π*(***θ***).

### 2.4 Bayesian adaptive estimation procedure

Given a randomly sampled observer, we estimate their representation by iterating three steps. Let *π*^(0)^(***θ***) = *π*(***θ***) and let *π*^(*i*)^(***θ***) denote the posterior after *i* iterations. At iteration *i*:

1. **Optimize distortion**. Compute a distortion direction ***ε***^(*i*)^ at a base stimulus ***s*** by optimizing Equation 2 with *π*^(*i*−1)^(***θ***) in place of the prior.
2. **Measurement likelihood**. Present the base stimulus ***s*** distorted along the direction ***ε***^(*i*)^ and record the observer’s perceptual judgments. Compute the likelihood *ℓ*(data| ***θ***) for each observer given the sampled observer’s perceptual judgments.
3. **Update posterior**. Compute the posterior using Bayes’ rule:

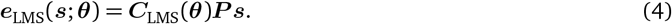

The specific measurement procedure and likelihood function depend on the application. We make each of these concrete in the next section.

### 2.5 Application: estimating an observer’s cone fundamentals

Color vision varies across individuals, even within color-normal populations, due to physiological differences in lens density, macular pigment, photopigment densities, and photopigment spectral absorbances which influence an observer’s CFs (CIE, 1989; Allen, 1970; Nayatani, Takahama, & Sobagaki, 1983; Ohta, 1985; Stiles & Burch, 1959; Emery & Webster, 2019; Bosten, 2022). We apply the adaptive estimation framework from the previous section to the problem of inferring a simulated observer’s CFs from perceptual discrimination judgments. The application has three components, described below: the 16-channel display of Oh et al. (2022) that defines the stimulus space, the individual colorimetric observer model of Asano, Fairchild, and Blondé (2016) that maps stimuli to internal representations and defines the prior distribution over observers, and the 3AFC oddity task used to measure an observer’s perceptual sensitivity. We report simulated experiments using this setup in the following section.

#### 2.5.1 Stimulus space defined by a 16-channel display

On an RGB display, a stimulus is specified by three channel intensities and the mapping to cone responses is generally modeled as a 3 × 3 linear transformation that is full-rank for any color-normal observer. While interobserver variability among color-normal observers leads to subtle differences in this transformation, the resulting differences in their color discrimination thresholds can be small and difficult to characterize within the gamut of a typical RGB display. A display with more spectral channels provides additional degrees of freedom that can be exploited to probe individual differences. We therefore model stimuli using the measured primaries of the 16-channel display described by Oh et al. (2022). Quantizing the visible spectrum into *w* discrete wavelengths (range: 400–700 nm; discretized step: 10 nm), the display’s channels can be described by a *w* ×16 matrix ***P*** that stacks the 16 quantized channel primaries (Figure 2A). Given a stimulus vector of channel intensities ***s*** ∈ [0, 1]^16^, the emitted spectral energy is ***P***_***s***_ ∈ ℝ^*w*^.

**Figure 2:**
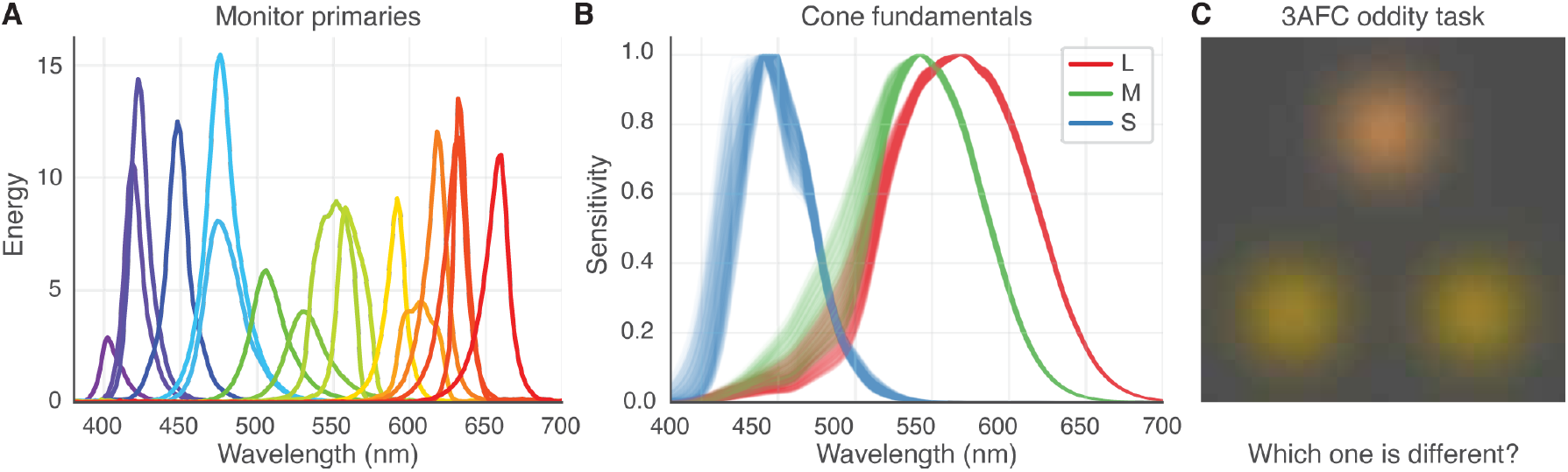
Set up for simulations estimating individual CFs. **A.** Spectrum of each of the monitor’s 16 channels. **B**. 1000 CFs sampled from the observer model. **C**. Schematic illustrating the 3AFC oddity task. Three stimuli are presented—two of them are identical with the location of the different stimulus randomized on each trial—and the observer’s task is to identify the odd one out. Note that the spatial configuration shown here is illustrative and does not play a role in the simulated experiments.

#### 2.5.2 Individual colorimetric observer model and stimulus encoding

We model individual variability in CFs within color-normal populations using the individual colorimetric model of Asano, Fairchild, and Blondé (2016). This model takes age and visual field as known covariates and decomposes the remaining variability in CFs into eight biophysical parameters: lens pigment density, macular pigment density, the optical densities of the L, M, and S photopigments, and shifts in the peak spectral sensitivities of the L, M, and S cones. Each biophysical parameter is expressed as a percentage deviation from its population mean, with standard deviations fit to empirical data. These parameters are modeled as independent and normally distributed. Together, these parameters determine an individual’s CFs, which can be described by a 3 × *w* matrix ***C***_LMS_(***θ***). The cone responses to a stimulus ***s*** ∈ [0, 1]^16^ on the 16-channel display are then

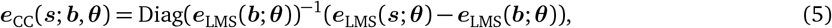

The CFs and their variability across all eight biophysical parameters are shown in Figure 2B.

Given the cone responses, we model the observer’s internal stochastic representation following Hong, Chow, Guan, Brainard, and Williams (2026). A stimulus ***s*** presented against a fixed background ***b*** ∈ [0, 1]^16^ is encoded as cone contrast:

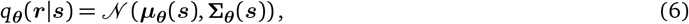

which normalizes each cone’s response by its adaptation level to the background. The representation is then a 3-dimensional Gaussian:

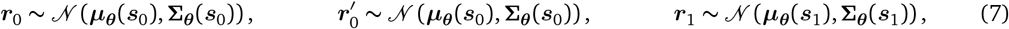

where ***µ***_***θ***_ (***s***) is an affine transformation of ***e***_CC_(***s***; ***b, θ***) and ***Σ***_***θ***_ (***s***) is the covariance field that captures internal noise in color perception (see section *Observer model* of the Supplement). The covariance field is constrained by approximately 30,000 3AFC color discrimination judgments from a single participant collected in a standard three-channel RGB display (Hong et al., 2026). These data are fit with a model that assumes the internal noise varies smoothly across stimulus space, so that small changes in the stimulus produce only small changes in internal noise. Once fit, the model yields the internal noise at any location within the RGB gamut. In the present simulations, however, stimuli are specified by the intensities of the 16 display channels. To assign an internal-noise covariance matrix to a synthetic observer at a given stimulus, we first compute the RGB values that would produce the same cone contrast in the reference human observer as in the synthetic observer, read out the covariance matrix at that RGB location, and use it as the synthetic observer’s internal noise.

#### 2.5.3 Color discrimination task

We simulate observers’ discrimination performance on a 3AFC oddity task (Figure 2C). On each trial, the simulated observer is presented two identical reference stimuli at channel intensities ***s***_0_ and one comparison stimulus ***s***_1_ = ***s***_0_ + Δ***ε***, presented against the fixed background ***b***, and identifies the odd one out. Here ***ε*** is the distortion direction and Δ is the perturbation weight (which can be positive or negative). For best results under gamut constraints, the reference stimulus ***s***_0_ is chosen based on the distortion ***ε*** (see section *Background and reference stimuli* of the Supplement).

From Equation 6, the internal representations of the three stimuli are given by

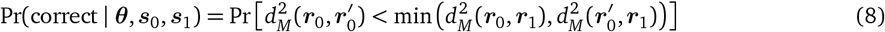

where ***r***_0_, 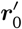 are the representations of the two reference stimuli and ***r***_1_ is the representation of the comparison (see section *Computing model-optimized stimulus distortions* of the Supplement). We assume these responses are independent and model the observer’s decision based on the Mahalanobis distances between pairs of internal representations: the two stimuli with the smallest pairwise distance are identified as the references, and the remaining stimulus as the comparison. The probability of a correct response lies in the range 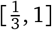 and is given by

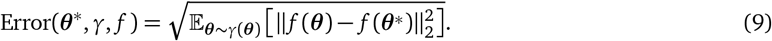

where 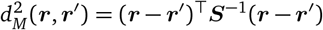and 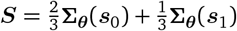 is the pooled covariance. Following Hong et al. (2025), we approximate Pr(correct | ***θ, s***_0_, ***s***_1_) via Monte Carlo simulation (2,000 samples).

To measure the observer’s discrimination performance at a base stimulus ***s***_0_ to a distortion direction ***ε***, we present repeated trials and vary Δ with a 2-down/1-up staircase targeting 70.7%. Because each channel of ***s***_0_+Δ***ε*** must remain within the display gamut [0, 1], we limit Δ so that the distorted stimulus remains in range. When the observer cannot reliably discriminate the perturbation even at the largest feasible weight (i.e., performance remains below 70.7%), we measure performance at that weight. Because the staircase varies the weight across trials, each trial *t* uses its own comparison ***s***_0_ + Δ_*t*_ ***ε*** with success probability *p*_*t*_ = Pr(correct| ***θ, s***_0_, ***s***_0_ + Δ_*t*_ ***ε***). Since the responses, *y*_*t*_ ∈ {0, 1}, are conditionally independent given the magnitudes, the likelihood is the product over trials:

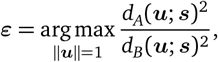

Evaluating this likelihood for each candidate observer ***θ*** yields the term used to update the posterior in Equation 3.

## 3 Results

We apply our adaptive estimation framework to synthetic observers generated from the individual colorimetric model of Asano, Fairchild, and Blondé (2016), using the 16-channel display of Oh et al. (2022), and the 3AFC task described above. For each experiment, we simulate the full adaptive procedure—distortion optimization, performance measurement, and posterior update—for a target observer drawn from the population. We use the fixed background across all experiments (see section *Background and reference stimuli* of the Supplement). To quantify the efficacy of our procedure, we define the error in estimating a function *f* (***θ***) of the true parameters ***θ***^∗^ under the distribution *γ*(***θ***):

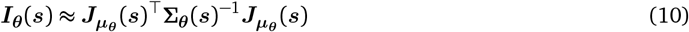

Here *f* (***θ***) is a function of interest such as the CFs, e.g. *f* (***θ***) = ***C***_LMS_(***θ***), or a biophysical parameter, e.g. *f* (***θ***) = *θ*_*k*_ for some *k*, and *γ*(***θ***) can be the prior distribution or a posterior distribution.

### 3.1 Experiment 1: Estimating an observer’s lens and macular density

We first consider the problem of estimating an observer’s lens and macular pigment densities. We draw a target observer (age 20) from the full eight-dimensional Asano population, but carry out inference over a model set that varies only lens and macular density, with the remaining six biophysical parameters assumed fixed at their population means. The target observer therefore does not in general lie within the model set, so the experiment also tests robustness to this mismatch. The model set is the two-dimensional grid of lens and macular densities shown in Figure 3A.

**Figure 3:**
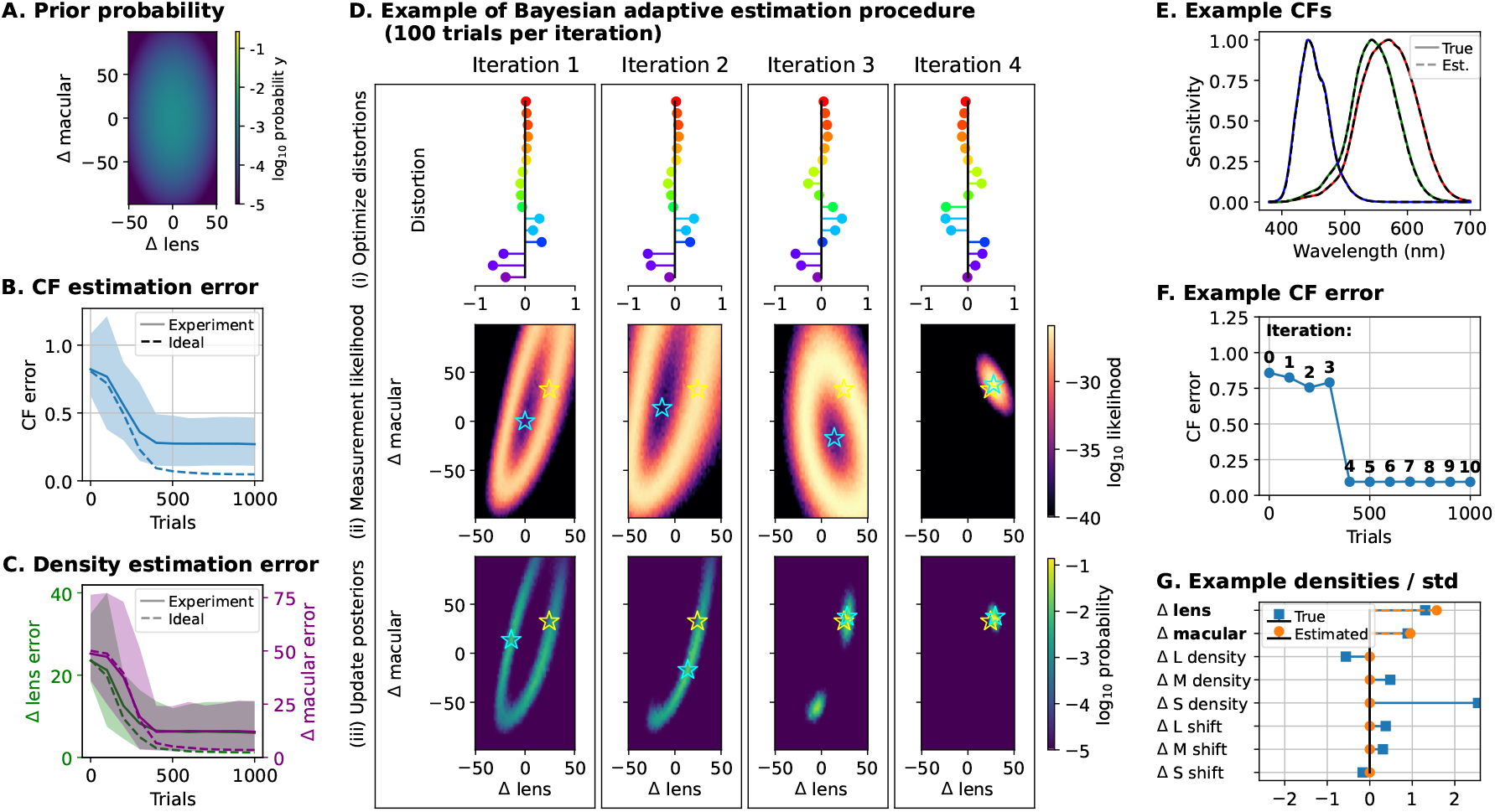
Experiment 1: estimating lens and macular density. Target observers are drawn from the full 8-dimensional population (aged 20), while inference is carried out over a 2-dimensional model set that varies only lens and macular density, with the other six parameters fixed at their population means. **A**. Prior over the 2D model set of lens and macular densities. **B**. CF estimation error versus trial number. Solid lines show the median and shaded regions the 5th–95th percentile range across 100 observers. **C**. Lens (left axis) and macular (right axis) density estimation error versus trial number, with median and 5th–95th percentile range as in B. In B and C, dashed lines show the median of an idealized procedure in which the six other biophysical parameters are fixed to their true values rather than the population means, which lower bounds performance achievable without access to the actual values. **D**. A single run of the adaptive procedure over five iterations (100 trials each): (i) the optimized distortion direction, (ii) the resulting per-observer likelihood, and (iii) the updated posterior, which concentrates around the target observer. **E**. Estimated (dashed) and ground-truth (solid) CFs for the example observer. **F**. CF estimation error versus trial number for the example observer. **G**. Estimated (dashed) and groundtruth (solid) Asano parameters (densities) divided by their standard deviations for the example observer.

We simulate the full adaptive procedure using batches of 100 simulated psychophysical trials per iteration. Across 100 sampled observers, both the CF error (Figure 3B) and the lens and macular density errors (Figure 3C) decrease over the first 500 trials before leveling off. Figures 3BC also show the performance of an “Ideal” algorithm in which lens density and macular pigment density are estimated but the other six biophysical parameters are fixed to their actual values rather than the population average. Note that in the “Ideal” case, the recovery of the lens and macular pigment densities is essentially perfect after 500 trials.

Figure 3D illustrates the adaptive procedure for a single observer drawn from the population model: at each iteration we optimize the distortion direction (row i), measure the observer’s performance and compute the per-observer likelihood (row ii), and update the posterior (row iii), which concentrates around the target observer across four iterations. The estimated CFs closely match ground truth for an example observer (Figure 3E). The CF error decays gradually across the first three iterations, drops significantly after the fourth iteration, and then levels off (Figure 3F). After four iterations, the estimated lens and macular densities are close to the true values, whereas the other parameters are fixed to their population averages (Figure 3G).

### 3.2 Experiment 2: Estimating all eight biophysical parameters

We next consider the general setting in which all eight biophysical parameters vary across observers. As before, the target observer is drawn from the full eight-dimensional model population at age 20, but inference is now carried out over an eight-dimensional model set spanning all eight parameters. Because an eight-dimensional grid would be prohibitively large, we instead construct the model set by clustering: we draw 12,000 observers i.i.d. from the prior and reduce them to 3,000 representatives over two rounds of agglomerative clustering, each round partitioning the models into pairs and replacing each pair with its centroid in parameter space. Because every group has the same size, each representative stands for the same number of prior samples, so the discrete set approximates the prior with uniform weights. Since the target observer is drawn from the continuous population, it will in general not coincide with any model in the set, introducing an additional source of estimation error.

Lens and macular density account for most of the variation in CFs and are the only parameters that are consistently estimable; the optical densities and peak-sensitivity shifts of the L, M, and S photopigments have weaker and partially overlapping effects on the CFs and remain poorly constrained over the course of the experiment (Figure 4A). This identifiability issue has been observed in other approaches to estimating CFs using color-matching procedures (Thomas & Mollon, 2004; Shi et al., 2024a). Nonetheless, because the estimable parameters are those that dominate CF variation, the procedure still recovers the observer’s CFs accurately: across 100 observers, the CF error decreases over trials even as most individual parameters remain uncertain. Figure 4B shows the best, median, and worst recoveries by CF error; even in the worst case the estimated CFs show only small deviations from the true CFs, while the per-parameter errors confirm that lens and macular density are recovered most reliably. In the next section, we compare the final performance of Experiment 1 (varying lens and macular density) and Experiment 2 (varying all eight biophysical parameters) for observers sampled from a range of ages.

**Figure 4:**
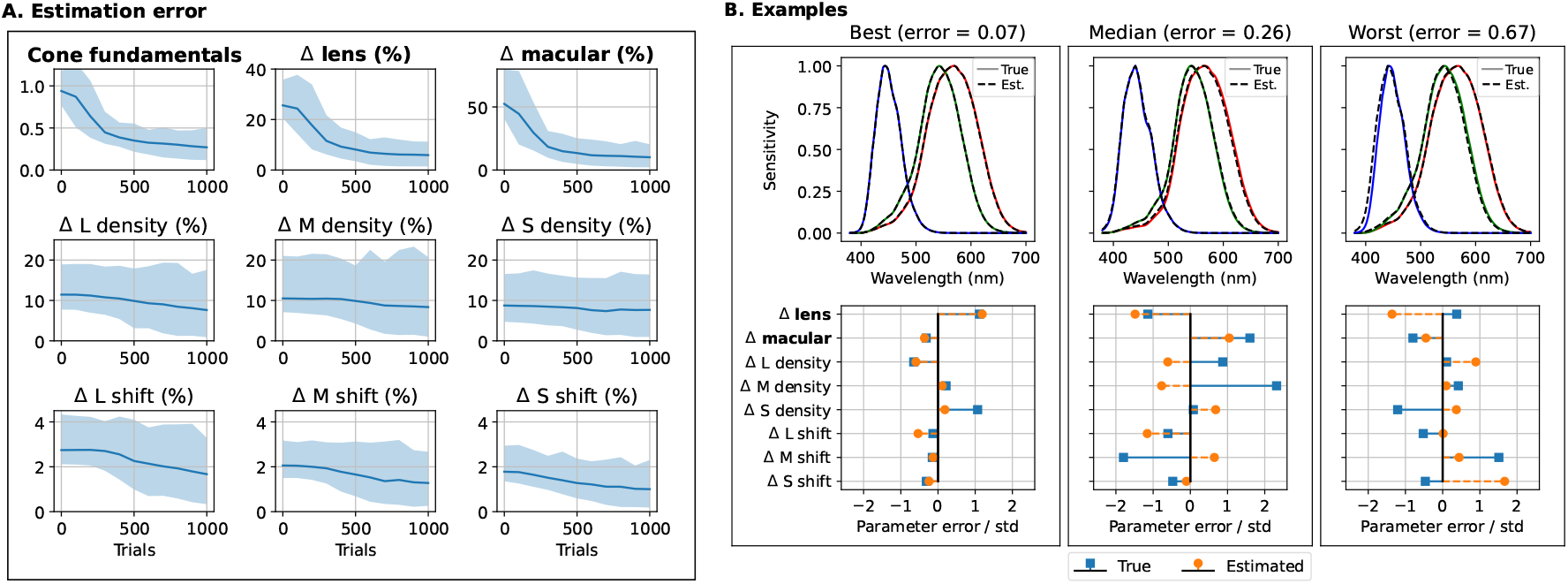
Experiment 2: estimating all eight biophysical parameters. Target observers are drawn from the full 8-dimensional Asano population (age=20), and inference is carried out over an 8-dimensional model set of 3,000 representatives constructed by clustering. **A**. Estimation error versus number of trials for the CFs and for each of the eight biophysical parameters; lines show the median and shaded regions the 5th–95th percentile range across 100 observers. **B**. Estimated (dashed) and ground-truth (solid) CFs (top) and per-parameter errors in units of standard deviations (bottom) for the best, median, and worst recoveries by CF error (error values in parentheticals).

### 3.3 Experiment 3: Estimation performance as a function of age

Finally, we examine how estimation performance depends on observer age and on the choice of model set. In the previous experiments age was fixed at 20; here we draw target observers across the age range 20–80 and treat age as a known covariate, asking how well the procedure recovers the remaining parameters as age varies. We also compare three model sets of decreasing richness: the 8-dimensional clustered set of Experiment 2, the 2-dimensional lens/macular grid of Experiment 1 (the other six parameters fixed at their means), and a 1-dimensional set varying only lens density (the other seven fixed at their means).

Figure 5 reports the final error after 1,000 trials as a function of age. The main result is that the 2D lens/macular model set recovers CFs about as well as the full 8D set, and substantially better than the 1D lens-only set, across the entire age range (Figure 5, top left). CF recovery is accurate and roughly stable with age for both the 2D and 8D sets, indicating that the procedure remains effective for older observers despite age-related changes in the lens. Estimation of lens density itself improves with age (Figure 5, top middle). By contrast, the 1D set yields substantially larger and more variable CF errors, confirming that lens density alone is insufficient to capture individual CFs. This is consistent with Experiment 2: lens and macular density dominate the variation in CFs and are the parameters the procedure estimates most reliably, so adding the remaining six parameters to the model set yields little further improvement in CF recovery.

**Figure 5:**
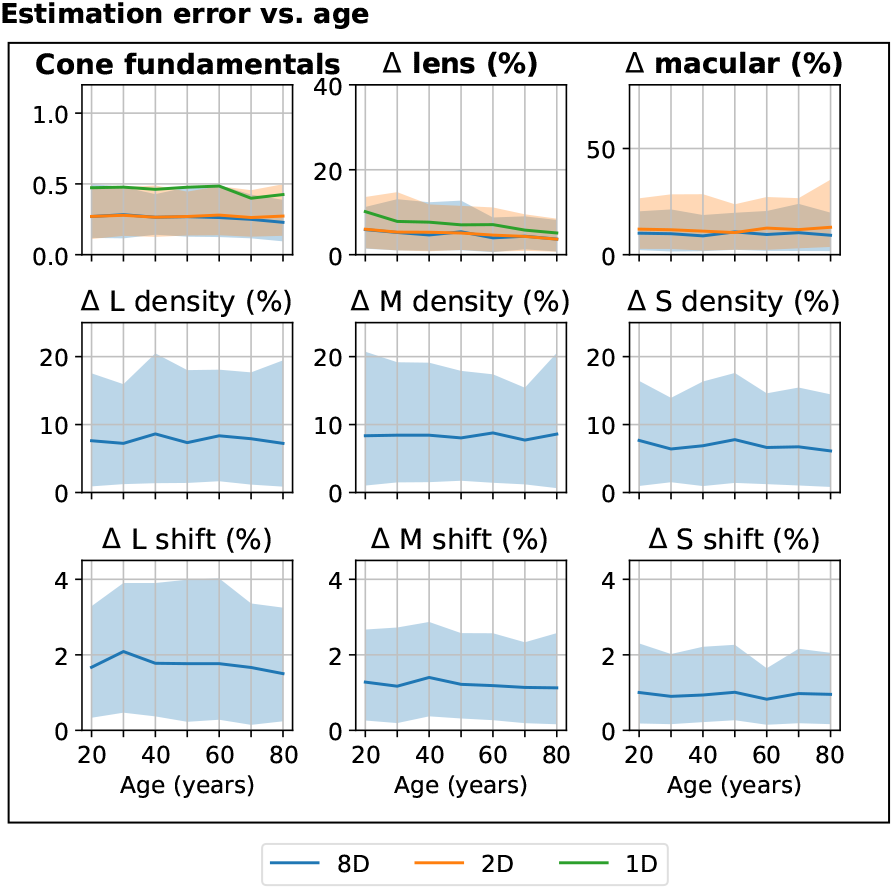
Experiment 3: comparing estimation performance versus age. Final estimation error after 1,000 trials for the CFs and each of the eight biophysical parameters, as a function of observer age. Target observers are drawn from the full 8-dimensional Asano population with age treated as a known covariate. Lines compare three model sets: 8D centroid models from clustering (as in Experiment 2), a 2D lens/macular grid (other six parameters fixed at their means, as in Experiment 1), and a 1D lens-only set (other seven parameters fixed at their means). Shaded regions show the 5th–95th percentile range across observers; the range for the 1D set is omitted for clarity, as it is much larger than the others.

## 4 Discussion

### 4.1 Comparison to other adaptive methods and model-comparison methods

Our method belongs to a broader family of adaptive and optimal experimental design approaches (MacKay, 1992; Chaloner & Verdinelli, 1995; Cohn, Ghahramani, & Jordan, 1996) and their application in psychology and neuroscience (Watson & Pelli, 1983; Watson, 2017; Benda, Gollisch, Machens, & Herz, 2007; Lesmes, Jeon, Lu, & Dosher, 2006; Lesmes, Lu, Baek, & Albright, 2010; DiMattina & Zhang, 2013; Pillow & Park, 2016; Bak & Pillow, 2018). It is most closely related to methods that generate stimuli to distinguish competing models (Wang & Simoncelli, 2008; Berardino et al., 2017; Golan et al., 2022; Zhou et al., 2024; Feather et al., 2025), which synthesize stimuli that maximally separate a fixed set of candidate models in order to decide which best accounts for an observer’s behavior. We build on the same FIM machinery as previous works (Berardino et al., 2017; Zhou et al., 2024; Feather et al., 2025), but our method is designed to exploit the smooth structure of a continuous population.

A further advantage concerns how informative stimuli are selected. General-purpose Bayesian adaptive methods such as QUEST+ (Watson, 2017) select each stimulus by minimizing the expected entropy of the posterior, evaluated over a discretized grid of candidate stimuli and parameters. This grid grows multiplicatively with the number of stimulus and parameter dimensions, so entropy minimization becomes intractable in high-dimensional stimulus spaces such as our 16-channel display. By contrast, our FIM-based criterion yields the maximally informative distortion analytically from the observer model, sidestepping the curse of dimensionality. This makes optimizing distortions tractable within a closed-loop adaptive procedure, where stimuli must be selected in real time between trials.

### 4.2 Comparison to other methods for estimating spectral sensitivities

Current methods for identifying individual differences in CFs of color normal individual typically use color-matching experiments, in which an observer adjusts a mixture of primaries until it matches a reference mixture (Stiles & Burch, 1959; Asano, Fairchild, Blondé, & Morvan, 2016). Recent work employs this approach with 13-primary LED instruments on 100 observers of varying age (Shi et al., 2024a, 2024b). For estimating properties of the L and M CFs in color normals, color matches restricted to longer wavelength stimuli (Rayleigh matches) have been also employed (Neitz & Jacobs, 1986, 1990; He & Shevell, 1994; Sanocki, Shevell, & Winderickx, 1994; Jordan & Mollon, 1995). There are two main differences between these works and ours. First is the perceptual experiment: our work uses perceptual discrimination judgments, whereas those works use color-matching experiments. Second is the stimulus selection process: the color-matching works present the same stimuli to every observer and then estimates the observer’s parameters, whereas our procedure adaptively select the distortions that best differentiate observers under the current posterior. Both approaches are subject to challenges in uniquely identifying underlying biophysical parameters (Shi et al., 2024a; Thomas & Mollon, 2004).

A related estimation problem arises in camera calibration, where the goal is to recover the spectral sensitivities of a camera without a monochromator (Walowit, Buhr, & Wüller, 2017). These methods exploit the fact that camera spectral sensitivities vary along only a few dimensions across a population of devices (Jiang, Liu, Gu, & Süsstrunk, 2013). This parallels our setting, in which individual variation in CFs is low-dimensional and an observer’s CFs are recovered as coordinates within a population model. The differences are the source of information and the choice of stimuli: these methods recover sensitivities from a device’s physical responses to a fixed set of characterization spectra, whereas ours infers them from an observer’s perceptual judgments to adaptively chosen distortion directions.

We also tested whether CFs could be recovered directly in their top principal components, rather than through the biophysical parameters. We compared estimating all eight model parameters against estimating CFs directly within their top three principal components, and found that the parameter-based approach recovered CFs more accurately (not reported). Both approaches reduce the effective dimensionality of the estimation problem: the principal-component approach does so by linearly projecting the CFs, whereas the parameter-based approach does so through a model-reduction step that clusters a large sample of observers into a representative finite set. The difference is that clustering preserves the nonlinear mapping from biophysical parameters to CFs, whereas a linear principal-component parameterization does not.

### 4.3 Future directions

Our results are a proof of concept on simulated observers, and the natural next step is to run the procedure on human observers using the 16-channel display of Oh et al. (2022). Such experiments would also test our assumption that the internal-noise field estimated from a single participant on a three-channel display (Hong et al., 2026) transfers to new observers viewing stimuli on the 16-channel display.

Beyond CFs, our framework requires only a parametric model of how an individual’s sensory representation varies within a population. It could therefore be applied to other sources of individual variation for which such computational models exist, such as variability in retinal ganglion cell responses (Shah et al., 2022).

## 5 Conclusions

We introduced a Bayesian adaptive procedure for estimating an individual observer’s sensory representation from targeted perceptual discrimination judgments. A key component of our approach is the selection of stimulus distortions based on a parametric model of observer variability in a population: at each iteration we choose a distortion that the modal observer is insensitive to relative to the population average. The optimal distortion is the solution to a generalized eigenvalue problem so it can be efficiently computed using standard spectral methods, making the optimization fast enough for a real-time closed loop experiment.

We applied our method to the problem of estimating individual CFs, using the individual colorimetric model of (Asano, Fairchild, & Blondé, 2016), the 16-channel monitor of (Oh et al., 2022), and 3AFC oddity judgments. The procedure recovered the CFs, as well as the lens and macular pigment densities, of simulated observers using approximately 500 3AFC oddity judgments, which corresponds to an experimental time of one hour. Performance was comparable whether we estimated all eight biophysical parameters or only the two governing lens and macular pigment densities, so restricting the observer model to these two parameters could reduce the procedure’s computational cost with little loss in accuracy.

## 6 Supplementary material

### 6.1 Observer model

Our observer model (Equation 6) requires specifying, for each synthetic observer ***θ***, both the mean ***µ***_***θ***_ (***s***) and covariance ***Σ***_***θ***_ (***s***) of the internal representation at any stimulus ***s***. The mean is determined by the observer’s cone contrast response, which depends on their cone fundamentals ***C***_LMS_(***θ***). The covariance is provided by the WPPM of Hong et al. (2025), which specifies a smoothly varying field of covariance matrices across stimulus space.

A complication arises because the WPPM was fit to participants’ data collected on a standard 3-channel RGB display. It defines a covariance field ***Σ***_WPPM_(·) over a three-dimensional *model space*—an affine transformation of the RGB coordinates of that display, bounded between −1 and 1 along each axis (Hong et al., 2025). Our synthetic observers, by contrast, have varying cone fundamentals and view stimuli on a 16-channel display. To bridge this gap, we adopt the following assumption: *two observers who experience the same cone contrast response to a stimulus share the same internal noise*. Under this assumption, we can query the WPPM for any synthetic observer by finding the location in model space that corresponds to their cone contrast response.

Given a synthetic observer ***θ*** viewing a stimulus ***s*** against background ***b*** on the 16-channel display, we proceed as follows.

**Step 1: Cone contrast**. Compute the observer’s cone excitations ***e***_LMS_(***s***; ***θ***) and cone contrast responses ***e***_CC_(***s***; ***b, θ***) using Equations 4–5.

**Step 2: Cone-contrast matching**. Identify the stimulus on the reference observer’s 3-channel RGB display that would produce the same cone contrast. Denoting the reference observer’s cone fundamentals as 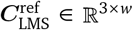 and their display’s primaries as ***P*** _RGB_ ∈ ℝ ^*w* × 3^, the reference observer’s cone excitations to the mid-gray background 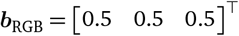 are 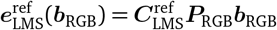. The cone excitations that would produce the matching cone contrast are:

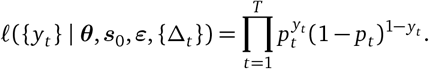

where 1 = [1 1 1] ^T^ is the vector of ones, and the corresponding RGB coordinates are recovered via the pseudo-inverse:

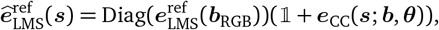

**Step 3: Query the WPPM**. The WPPM’s model space is related to RGB by the affine mapping ***s***_RGB_ ↦ 2***s***_RGB_ − 1 . We form the internal representations by querying the WPPM at the matched model-space coordinates:

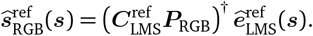

where

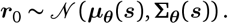

### 6.2 Computing model-optimized stimulus distortions

This section makes the modal-observer objective of Equation 2 concrete for the individual colorimetric model, yielding a closed-form expression for the optimized distortion direction.

Using the facts that ***r*** is normally distribution and **Σ**_***θ***_ (***s***) varies slowly with ***s***, we approximate the FIM as

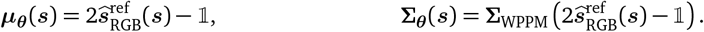

where

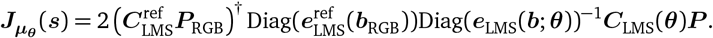

is the Jacobian of ***µ***_***θ***_ (***s***) with respect to ***s***.

Recall that Equation 2 selects the distortion to which the modal observer, ***θ***_modal_ = arg max_***θ***_ *π*(***θ***), is least sensitive relative to the population average. In the colorimetric setting, the modal observer’s cone responses depend on the stimulus only through ***C***_LMS_(***θ***_modal_)***P*** ∈ ℝ^3×16^(Equation 4), so any stimulus direction in the null space of this map leaves the modal observer’s representation unchanged. Its Fisher information 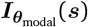 is therefore rank-deficient, with a 13-dimensional null space to which the modal observer is insensitive. We collect an orthonormal basis for this null space in the columns of ***K*** ∈ ℝ^16×13^, i.e., the columns of ***K*** are orthonormal and span the null space { ***s*** : ***C***_LMS_(***θ***_modal_)***P s*** = **0** .}

Because the denominator of Equation 2 vanishes on ***K*** while the numerator remains positive, the ratio is unbounded there, so the optimum lies in span(***K***). Among those directions we select the one that maximizes the numerator, the posterior-weighted expected sensitivity across the population:

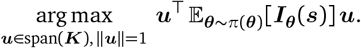

Intuitively, because the observer model varies smoothly with ***θ***, models near the modal observer in ***θ***-space are also nearly insensitive to distortions in span(***K***) and contribute little to this expectation. The population-averaged objective is therefore likely dominated by models far from the mode.

We use the FIM approximation from Equation 10 and write ***u*** = ***Kx*** with ***x*** ∈ ℝ^13^ to rewrite the objective as follows (suppressing the dependence on ***s***):

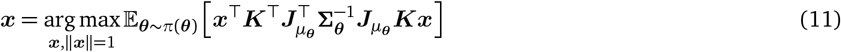

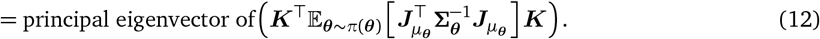

The optimized distortion direction is then recovered as ***ε*** = ***Kx***. In practice, we calculate ***Σ***_***θ***_ around ***s***_0_ = [0.5]^16^.

### 6.3 Background and reference stimuli

As described in Color discrimination task, in each trial the simulated observer is presented two identical reference stimuli ***s***_0_ and a perturbed version ***s***_1_ = ***s***_0_ + Δ***ε***, all against the fixed background ***b***, and identifies the odd one out. We use the following fixed background stimulus ***b*** across all experiments:

[0.4764, 0.5025, 0.4846, 0.5379, 0.6667, 0.6424, 0.6753, 0.6856, 0.6071, 0.5378, 0.4763, 0.4053, 0.3594, 0.3771, 0.4039, 0.3555], chosen to match the background used by Hong et al. (2026), whose data constrain our internal-noise model.

Given the fixed background ***b***, we aim to make candidate observers as distinguishable as possible, which depends on the reference stimulus ***s***_0_, the distortion direction ***ε*** and the distortion magnitude Δ. We proposed a principled, closed-form method for choosing the distortion direction (Equation 2) and we adaptively choose Δ using a staircase procedure. We now explain our choice of reference stimulus.

An important factor when choosing the reference stimulus is the limited gamut of the monitor. In simulations, we often found that for a given ***s***_0_ and ***ε***, Pr(correct| ***θ, s***_0_, ***s***_0_ + Δ***ε***) stayed near chance for many observers even when Δ was pushed to the edge of the gamut, so that those observers could not be told apart at any admissible perturbation weight. Our optimization method does not capture this constraint. Accounting for it would require evaluating Pr(correct | ***θ, s***_0_, ***s***_0_ + Δ***ε***) by Monte Carlo simulation for every candidate observer and reference stimulus, adding to runtime cost. We instead compare a small set of candidate references and keep the one that best separates observers.

We quantify how well a distortion–reference pair separates observers by the expected pairwise distance between candidate observers’ accuracies, evaluated at the strongest perturbation weight the gamut allows:

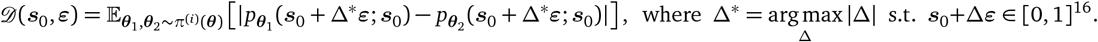

We compute the distortion direction ***ε*** from two variants of Equation 2, pair each with a small set of distortion-specific candidate references (***s***_0_, ***ε***), and keep the distortion-reference pair that maximizes . The two distortion variants and the candidate references are as follows.

Our two possible choices for distortion direction ***ε*** are:

1. *ε* from Equation 12, where ***Σ***_***θ***_ is calculated around ***s***_0_ = [0.5]^16^.
2. *ε* from Equation 12, where we fix the noise covariance ***Σ***_***θ***_ = ***I***.

Our possible choices for reference ***s***_0_ are:

1. Uniform: ***s***_0_ = [0.5]^16^. If we had to choose a single reference for all trials, this would be a very reasonable choice since our distortions often have both positive and negative channel intensities, so setting every channel towards the middle of its output range (i.e., 0.5) would allow each channel to increase or decrease as prescribed by the distortion. The strongest perturbation weight Δ^∗^ for a distortion ***ε*** in this case is 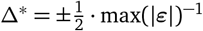
2. Sign-based: 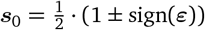. These distortion-dependent binary references (***s***_0_ ∈ {0, 1}^16^) increase the strongest perturbation weight to Δ ^*^ = ∓ (max(|***ε***|))^− 1^. When Δ goes from 0 to Δ^∗^, the channel corresponding to max(|***ε***|) will either go from 0 to 1 or 1 to 0 depending on the sign used above.
3. Scaled and sign-based: 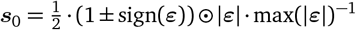. These distortion-dependent references have the same maximum perturbation weight Δ^∗^ = ∓ (max( |***ε***| ))^−1^ as the previous sign-based references, but also minimize the magnitude of the reference: each channel is anchored at 0 or at| ***ε***_*k*_| max( |***ε***|)^−1^ rather than at 0 or 1. As Δ goes from 0 to Δ^∗^, channel *k* sweeps between 0 and |***ε***_*k*_| max( |***ε***| )^−1^, so the channel carrying the largest |***ε***_*k*_ |spans the full [0, 1] range while every channel reaches the 0 boundary at one end of its sweep, using as much of the display’s dynamic range as the gamut permits.

## Notes

### Competing Interest Statement

The authors have declared no competing interest.

